# Autistic Traits are Associated with Suboptimal Decision Bias Strategies in Subsecond Timing

**DOI:** 10.64898/2026.05.11.724252

**Authors:** Matteo Frisoni, Luca Tarasi, Sara Borgomaneri, Vincenzo Romei

## Abstract

Time perception difficulties are frequently reported in Autism Spectrum Disorder, yet empirical findings remain inconsistent. A key methodological limitation is the failure to separate perceptual sensitivity from decision-making strategies. We applied Signal Detection Theory (SDT) to a subsecond duration discrimination task (100 and 500 ms) in 65 non-clinical adults varying in autistic traits, assessed via the Autism-Spectrum Quotient (AQ) and a Principal Component Analysis (PCA) of its subscales. Autistic traits did not predict reduced perceptual sensitivity (d′): temporal discrimination remained intact across the full autism-trait continuum, with Bayesian analyses providing converging evidence against a perceptual deficit. Instead, a PCA-derived cognitive component — combining heightened Attention to Detail with reduced Imagination — was systematically associated with a shift in decision bias (c). Individuals with this profile showed a graded attenuation of standard-based anchoring, with ordinal position progressively filling the gap. This shift operated consistently across both temporal scales, as confirmed by trial-level generalized linear mixed modelling, and reflects a quantitative redistribution of anchoring weight rather than a categorical switch in strategy. These findings reframe temporal “rigidity” in ASD not as a perceptual deficit, but as a suboptimal yet internally consistent decision-making style favouring within-trial information over accumulated representational knowledge.

**Lay Abstract:** Many autistic people report difficulties with time in daily life, but scientists have long disagreed on whether this reflects a genuine perceptual problem. This study found that autistic traits do not impair the basic ability to judge duration. Instead, people with more autistic traits tend to rely on which event came first, rather than accumulating experience across trials to refine their judgments — a less effective but internally consistent strategy.

## Introduction

Autism Spectrum Disorder (ASD) is a neurodevelopmental condition characterized by deficits in social interaction and communication, as well as restricted and repetitive behaviors and interests (DSM-5; APA, 2013). Beyond these diagnostic features, individuals with ASD frequently report difficulties managing the temporal structure of everyday life — anticipating events, adapting to schedules, and coping with transitions. Time perception, broadly defined as the ability to estimate and compare durations (Block, Grondin, & Zakay, 2018), underlies planning, episodic memory, and predictive behavior. Yet the nature of temporal processing in autism remains debated (Allman & Falter, 2015; Casassus et al., 2019).

Empirical findings are inconsistent. While some studies report intact or superior subsecond discrimination in ASD (Mostofsky et al., 2000; Falter et al., 2012, 2013), others find elevated thresholds or increased variability (Grondin, 2020; Bhatara et al., 2013; Kargas et al., 2015). A key methodological limitation is the reliance on measures such as the Point-of-Subjective-Equality or discrimination thresholds that cannot distinguish perceptual sensitivity from systematic decision biases (Block et al., 2018). The possibility that variability instead lies in the strategy used to perform temporal comparisons has rarely been considered.

A powerful probe of this issue is the standard-position effect: sensitivity is reliably higher when a fixed standard interval appears first in a sequence than when it appears second (Hellström & Rammsayer, 2015; Dyjas, Bausenhart, & Ulrich, 2012; Lapid et al., 2008). This effect reflects the dynamic construction of an internal reference across trials (Internal Reference Model; Dyjas et al., 2012; Lapid et al., 2008), in which repeated exposure to an identical standard stabilizes a representational trace that anchors subsequent comparisons. Constructing such a model is a prior-building operation requiring the integration of experience across trials.

Theoretical accounts of autism converge on precisely the type of representational difference that would predict a shift from standard-based toward positional anchoring. Predictive coding frameworks (Pellicano & Burr, 2012; Van de Cruys et al., 2014) propose attenuated top-down priors relative to bottom-up signals (Ursino et al., 2022; Tarasi et al., 2022). Weak central coherence (Happè & Frith, 2006) describes reduced integration of information into global representations. The enhanced perceptual functioning account (Mottron et al., 2006) describes heightened sensitivity to immediate input with reduced contextual integration. If cross-trial prior-building is less efficiently engaged along the autism-trait continuum, the consequence should not be impaired duration judgment, but a strategic shift in anchoring: less reliance on the stable standard-based reference, more on simple ordinal position.

In this context, two dimensions of autistic traits, as measured by the Autism-Spectrum Quotient (AQ; Baron-Cohen et al., 2001), are particularly relevant. The Imagination subscale indexes a reduced tendency to generate flexible internal representations and to simulate non-present events mentally. These are capacities that plausibly support the construction and maintenance of an internal model of the standard across trials. The Attention to Detail subscale indexes processing of individual features in isolation rather than integration across instances. Together, this cognitive profile characterizes a system reactive to immediate sensory input but less inclined to build accumulated representational knowledge.

Signal detection theory (SDT; Green & Swets, 1966; Romei & Tarasi, 2026) enables the separation of sensitivity and response bias. Although SDT is widely used in psychophysics, SDT has rarely been applied to temporal perception (Block et al., 2018; Frisoni et al., 2025), despite its potential to help distinguish perceptual differences from decisional strategies. Two conceptually distinct forms of bias can be separated: a standard-bias criterion, indexing anchoring to the repeated standard regardless of ordinal position; and an order-bias criterion, indexing preference for the first versus second stimulus independent of standard identity, i.e., the classical time-order error (Fechner, 1860; Hellström & Rammsayer, 2015). If cross-trial prior-building is less efficiently engaged along the autism-trait continuum, individuals higher in autistic traits should rely less on the adaptive standard-based anchor and more on the simpler positional heuristic — manifesting as stronger order bias without necessarily impairing perceptual sensitivity.

We applied SDT to a subsecond duration discrimination task (100 vs. 500 ms) to test whether autistic traits predict reduced temporal sensitivity or systematic decision biases. We supplemented this with a principal component analysis (PCA) of the five AQ subscales, designed to identify latent components reflecting cognitive styles characterized by heightened attention to detail and reduced imagination (Happè & Frith, 2006; Baron-Cohen, Ashwin, Aswhin, Tavassoli, Chakrabarti, 2009). We hypothesized that autistic traits would not predict reduced sensitivity, but a graded attenuation of standard-based anchoring with ordinal position filling the gap, a quantitative redistribution of decisional weight rather than a categorical switch in strategy.

## Methods

### Participants

Sixty-five participants (31 male; aged 19–31 years) were included after excluding two outliers whose accuracy scores fell more than two standard deviations from the group mean on both temporal discrimination tasks. Sample size was determined following recommendations from Samaha and Romei (2024). All participants provided written informed consent.

### Procedure

Two duration discrimination tasks were administered in a two-interval forced-choice (2IFC) paradigm (Figure 1). The Short Task used a 100 ms standard; the Long Task used a 500 ms standard. These durations were selected to span both minimal and longer sub-second intervals. Task order was counterbalanced across participants. On each trial, two checkerboard stimuli were presented sequentially ; participants indicated which lasted longer via keyboard press (“m” = first stimulus; “k” = second). A fixed contrast level (RGB 35/220; Tarasi & Romei, 2024) was used while varying the duration of stimulus presentation.

**Fig. 1.**
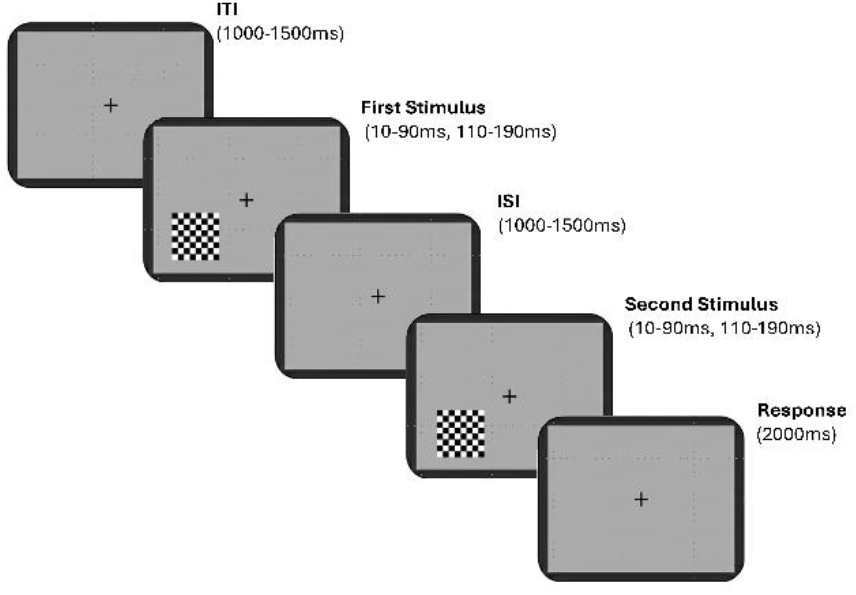
Experimental paradigm for the Short Task (100 ms) Two checkerboards were presented sequentially, separated by a fixation cross; participants judged which lasted longer, responding within 2 s (“m” = first; “k” = second). The Long Task (500 ms) was identical except for stimulus duration.

The standard stimulus was constant across trials; the comparison varied from ±10% to ±90% of the standard (method of comparison; Grondin, 2010). Standard and comparison positions were randomized and balanced (roving method; Macmillan & Creelman, 1991). Participants completed 360 trials per task (720 total), with 20 trials for each ratio level. Each trial included: fixation cross (1000–1500ms jitter), first stimulus, ISI (1000–1500ms jitter), second stimulus, response window (2000ms). The experiment was completed in a single session with three short breaks.

### Autistic traits

All participants completed the Autism-Spectrum Quotient test (AQ; Baron-Cohen et al., 2001), a 50-item self-report measure comprising five 10-item subscales: Imagination (assesses imaginative ability), Communication (assessing the weakness in communication skills), Social Skills (assessing the presence of poor social skills), Attention to Detail (assessing the exceptional attention to detail), and Attention Switching (assessing poor attention switching ability/strong focus of attention).

Items are scored dichotomously (0/1); total scores range from 0 to 50 (population mean ≈ 17; Ruzich et al., 2015). The Italian version was used (Ruta et al., 2018). A principal component analysis (PCA) with Varimax rotation was conducted on the five subscales to identify latent cognitive components (see below).

### Data Analysis

We applied SDT (MacMillan & Creelman, 1991) to each participant’s responses. The signal condition corresponded to trials in which the standard was longer; the noise condition to trials in which the comparison was longer. Responses were coded as Hits, Misses, False Alarms, or Correct Rejections, allowing computation of sensitivity (*d*′), which quantifies discrimination independently of bias, and criterion (*c*), which captures systematic tendencies in choice behavior.

We computed two criterion indices: the standard-bias criterion (Hellström & Rammsayer, 2015), where negative values indicate a bias toward the standard and positive values toward the comparison; and the order-bias criterion (Time Order Error; Fechner, 1860), where negative and positive values indicate preferences for the first and second-presented stimulus, respectively. Individual differences in autistic traits were assessed via the AQ (see *Autistic traits* section). To capture latent patterns beyond the total score, we conducted a Principal Component Analysis on the five subscales to identify cognitive styles potentially explaining the results better than the overall sum.

Statistical analyses proceeded in two stages. The first stage characterized sample-level performance. We conducted 2 × 2 within-subjects ANOVAs on d’ and criterion with TASK (Short vs. Long) and STANDARD POSITION (First vs. Second) as within-subjects factors. We then examined whether autistic traits predicted perceptual sensitivity using repeated-measures ANCOVAs with AQ total score and the PCA-derived component as covariates, supplemented by Spearman correlations and Bayesian Pearson correlations. Bayesian model comparison evaluated whether including AQ or the PCA component improved model fit for d′ relative to task and position alone.

The second stage examined whether autistic traits predict decision bias. The primary analysis was a trial-level Generalized Linear Mixed Model (GLMM) with a binomial family and logit link function. The dependent variable was ChoiceFirst, a binary indicator of whether the participant selected the first-presented stimulus on each trial (1 = first ; 0 = second stimulus chosen). Fixed effects included StandardPosition, PCA score, Task, and the interactions StandardPosition × PCA and Task × PCA; subject-specific random intercepts accounted for repeated measures. Fixed-effect significance was assessed using likelihood ratio tests (χ^2^). This trial-level approach leverages approximately 700 observations per participant and avoids the algebraic cancellation that would arise when collapsing across standard positions in SDT criterion analyses: because the standard-bias criterion reverses sign by construction across standard positions, any omnibus analysis would cancel two opposing effects, producing a spurious null result rather than a true absence of association.

To decompose the structure of the effect identified by the GLMM, we complemented it with condition-specific SDT criterion analyses. Spearman correlations and linear regressions were computed separately for each combination of task and standard position, preserving the conditional structure of the bias reversal.

As a complementary index of perceived duration, we computed individual Points-of-Subjective-Equality (PSE) from fitted psychometric functions. As PSE does not dissociate perceptual sensitivity from decision bias, full methodological details and results are reported in the Supplementary Materials.

## RESULTS

### 1. Sample-Level Discrimination Performance

#### Sensitivity (d’)

A 2×2 within-subjects ANOVA on d’ revealed three significant effects (Fig. 2A). The main effect of TASK was significant (F(1,64)=16.36, p<.001; η^2^p=.20), with higher sensitivity in the Long Task (1.60±0.49) than in the Short Task (1.43±0.44). The main effect of STANDARD POSITION was also significant (F(1,64)=155.48, *p*<.001; η^2^p=.71), with greater sensitivity when the standard appeared first, in line with the well-established standard-position effect (Hellström & Rammsayer, 2015). A significant TASK × STANDARD POSITION interaction was also observed (F(1,64)=6.34, *p*<.05; η^2^p =.09). Tukey post hoc tests confirmed that sensitivity was greater in the Long Task than in the Short Task for both standard-first (p < .0005) and standard-second (*p* < .05) conditions, and consistently higher in standard-first than standard-second trials within each task (Short Task: 1.71 ± 0.55 vs. 1.25 ± 0.46; Long Task: 2.00 ± 0.51 vs. 1.38 ± 0.59; both *p* < .0005).

**Fig. 2.**
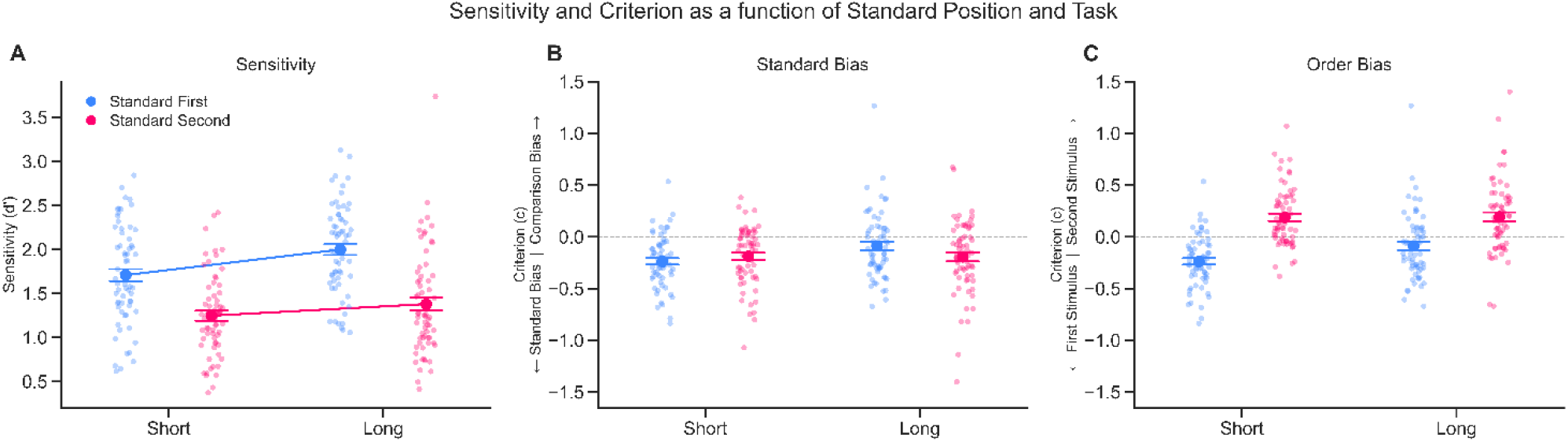
Sensitivity and biases across Task and Standard Position. In all panels, individual data points and mean ± SEM are shown. A. Sensitivity (d′): Sensitivity was higher when the standard appeared first (blue) than second (red), and in the Long Task relative to the Short Task, illustrating the standard-position effect and its interaction with task duration. **B. Standard Bias (criterion):** Negative values reflect a bias toward the standard stimulus, positive values toward the comparison. Criterion was consistently negative across conditions, with stronger bias in the Short Task, particularly when the standard appeared first. **C. Order Bias (criterion):** Negative values indicate a bias toward the first-presented stimulus, positive toward the second. On average, responses were driven by anchoring to the standard rather than a fixed positional preference.

The lower sensitivity in the Short Task likely reflects task-specific factors rather than a fundamental perceptual limitation for brief intervals, as sub-second durations are thought to be processed more automatically than longer intervals (Mioni, Stablum, & Grondin, 2013; Lewis & Miall, 2003). The robust standard-position effect across both tasks confirms the paradigm’s validity.

#### Criterion: Standard Bias

A 2 × 2 within-subjects ANOVA on criterion revealed a significant main effect of TASK (F(1,64) = 14.57, *p* < .001; η^2^p = .19), with more negative values in the Short Task (−0.21 ± 0.12) than in the Long Task (−0.15 ± 0.11), indicating stronger bias toward the standard under higher task difficulty (Fig.2B). The main effect of STANDARD POSITION was not significant (F(1,64) = 0.2, *p* = .658; η^2^p = .003). However, a significant TASK × STANDARD POSITION interaction emerged (F(1,64) = 5.07, p < .05; η^2^p = .09), reflecting particularly strong standard bias in the Short Task when the standard appeared first (−0.24 ± 0.25) compared with the Long Task in the same condition (−0.09 ± 0.32; post hoc *p* < .05).

One-sample t-tests confirmed a systematic bias toward the standard across all conditions. Collapsing across positions, criterion values were significantly negative in both the Short Task (−0.21 ± 0.12; *t*(64) = −14.01, *p* < .001, Cohen’s *d* = −1.74) and the Long Task (−0.15 ± 0.11; t(64) = −10.95, p < .001, Cohen’s *d* = −1.36). Condition-specific analyses confirmed that standard bias was significant and consistent across all four conditions (Short Task standard-first: −0.24 ± 0.25, t(64) = −7.60, *p* < .0001, Cohen’s *d*=−0.94; standard-second: −0.19 ± 0.28, t(64)=−5.31, *p*<.0001, Cohen’s *d* =−0.66; Long Task standard-first:−0.09 ± 0.32, t(64)=−2.24, *p*=.029, Cohen’s *d*=−0.28; standard-second: −0.19 ± 0.35, t(64)=−4.39, *p*<.0001, Cohen’s *d*=−0.55).

In sum, standard bias was consistent across conditions and amplified by task difficulty and standard-first presentation in the Short Task. This pattern of preserved sensitivity coupled with systematic standard bias establishes the baseline against which individual differences in autistic traits are evaluated.

#### Criterion: Order Bias

To examine whether participants displayed a generic preference for the first or the second stimulus (order bias; Fechner, 1860; Djas,Bausenhart et al.,2012) irrespective of whether it was the standard or the comparison, we recoded the SDT criterion to reflect first-vs. second-choice bias, with negative values indicating a tendency toward the first-presented stimulus and positive values toward the second.

Collapsing across presentation orders, no overall order bias was observed in either task (Short: −0.01±0.24; *t*(64)=−0.21, *p*=.836, Cohen’s *d*=−0.03; Long:0.07 ± 0.31; *t*(64) = 1.70, *p* = .094, Cohen’s *d* = 0.21). When separated by presentation order, however, biases tracked the position of the standard: participants favored the first stimulus when the standard appeared first (Short Task: −0.24 ± 0.25, *t*(64) = −7.60, *p* < .001, Cohen’s *d* = −0.94; Long Task: −0.09 ± 0.32, *t*(64) = −2.24, *p* = .029, Cohen’s *d* = −0.28), and the second when the standard appeared second (Short Task: 0.19 ± 0.28, *t*(64)=5.31, *p*<.001, Cohen’s *d* =0.66; Long Task: 0.19 ± 0.35, *t*(64)= 4.40,*p*<.001, Cohen’s *d* = 0.55; Figure 2C). These findings confirm that the dominant bias was toward the standard, not toward any fixed ordinal position.

### 2. Individual Differences: Autistic Traits and Temporal Performance

#### Autistic Traits and PCA-derived component

To uncover latent cognitive styles underlying individual differences in autistic traits, we conducted a Principal Component Analysis (PCA) on the five AQ subscales (Varimax rotation), yelding two orthogonal components. The AQ captures subclinical autistic traits across domains related either to social functioning (i.e., Social Skills and Communication) or to non-social aspects of cognition (i.e., Imagination, Attention to Detail, and Attention Switching). Importantly, factor-analytic studies have consistently shown that the empirical structure of the AQ deviates from its original five-subscale design, with prior work identifying between two and five factors and reporting considerable variability in item clustering (Austin, 2005; Hoekstra et al., 2008; Lau et al., 2013).

In our data, the second component accounted for 25.1% of the total variance and was characterized by strong positive loadings on Imagination (0.87) and Attention to Detail (0.63; see Supplementary Materials). Higher scores on these subscales reflect, respectively, reduced imaginative ability and enhanced focus on local features. Although typically treated as distinct factors, their co-loading here suggests a cognitive style marked by systematic, detail-oriented processing coupled with reduced imaginative flexibility. This component was also significantly correlated with the AQ total score (r = 0.62, *p*<.0001), underscoring its relevance as an index of autistic traits.

#### Autistic Traits Do Not Predict Perceptual Sensitivity

A repeated-measures ANCOVA with AQ total score as a covariate confirmed that the primary ANOVA effects replicated unchanged. Critically, AQ did not significantly predict d’ (F(1,63) = 0.34, *p* = .564, η^2^p = .005), nor did it interact with TASK or STANDARD POSITION (all ps >.37). An identical ANCOVA substituting AQ with the PCA-derived component yielded the same pattern: The PCA component did not predict d’, F(1,63) = 2.48, *p* = .121, η^2^p = .04), nor did it interact with either factor (all *ps* > .16).

Spearman correlations confirmed this null pattern.AQ total scores showed consistently weak, non-significant associations with d’ across all conditions (Short Task: rs = .01–.10, all *ps* > .42; Long Task: rs = .06–.11, all *ps* > .37; Collapsed: rs = −.11–.09, all *ps* > .40; see Figure 3, left). The PCA component showed similarly weak associations (Short Task: rs = .10–.20, all *ps* > .11; Long Task: rs = .19–.23, all *ps* > .07; see Figure 3, right). One correlation reached nominal significance when collapsing across tasks (standard-first: r = .26, *p*=.04), but failed to replicate in either task individually (Short: r = .20, *p* = .11; Long: r = .19, *p*=.14), suggesting sampling variability across 18 comparisons.

**Figure 3.**
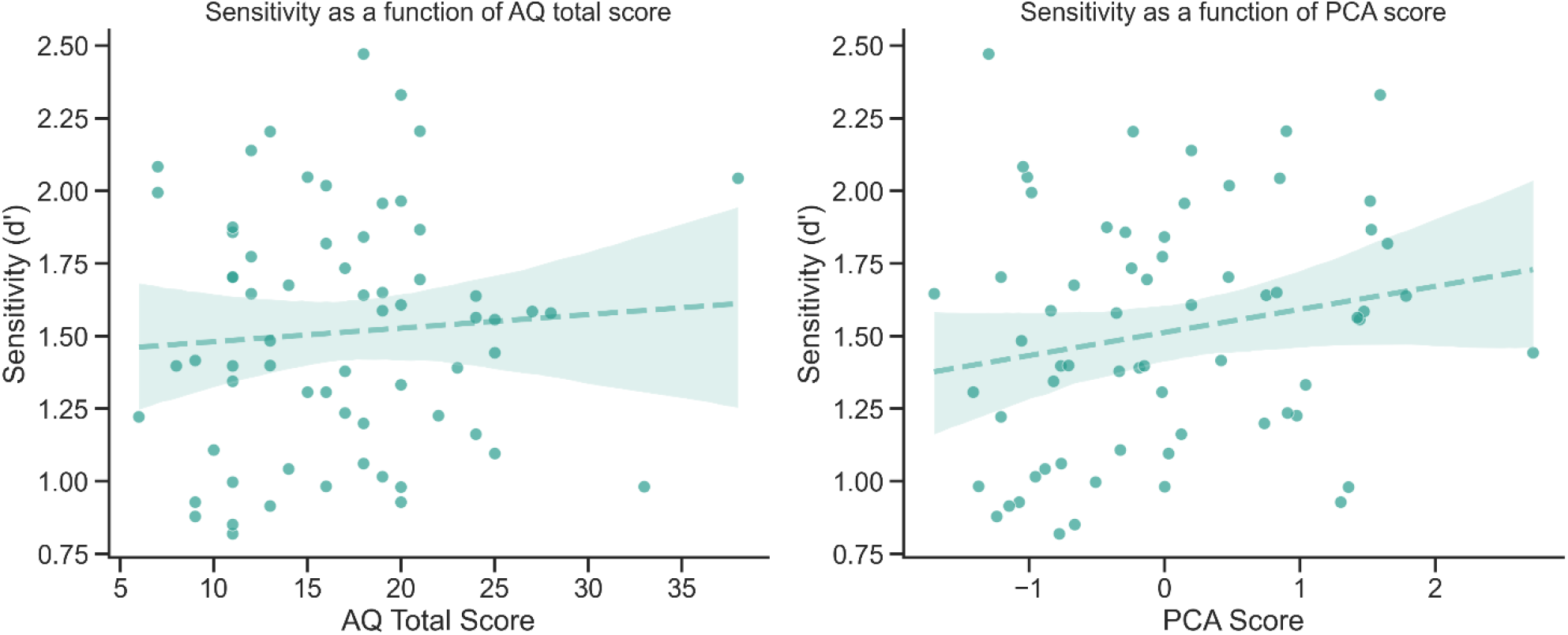
Sensitivity as a function of autistic traits, collapsed across tasks and standard positions. Each panel shows the relationship between individual *d*′ values (averaged across tasks and standard positions) and AQ total score (left) or PCA-derived component score (right), with regression lines and 95% CIs. Neither predictor was significantly associated with sensitivity (AQ: r = .07, BF_10_ =0.18; PCA: r=.19, BF_10_ =0.50). Both BF_10_ values below 1.0 indicate no support for an association with temporal sensitivity, consistent with condition-specific analyses. Note that the BF_10_ for the PCA component reflects inconclusive rather than strong evidence for the null, a pattern consistent with the small, unreliable trends observed across individual conditions.

Bayesian Pearson correlations provided converging evidence for the absence of a meaningful relationship between autistic traits and d’. In the Short Task, AQ–d’ correlations were negligible across all conditions (collapsed: r = 0.01, BF_10_ =0.16; standard-first: r=0.05, BF_10_ = 0.17; standard-second: r=−0.02, BF_10_ =0.16). The PCA component showed a comparable pattern (collapsed: r = 0.12, BF_10_ = 0.24; standard-first: r =0.16, BF_10_ =0.36; standard-second: r =0.09, BF_10_ =0.20). In the Long Task, AQ correlations remained weak (collapsed: r =0.12, BF_10_ =0.24; standard-first: r =0.06, BF_10_ =0.17; standard-second: r =0.13, BF_10_ = 0.26), as did PCA correlations (collapsed: r =0.23, BF_10_ = 0.84; standard-first: r =0.17, BF_10_ =0.37; standard-second: r =0.21, BF_10_ =0.58). Across both tasks collapsed, all Bayes Factors remained below 1.0, providing no support for the alternative hypothesis.

Bayesian model comparison reinforced this conclusion. The best-supported model included only TASK and STANDARD POSITION (P(M|data) = .386), while models incorporating AQ received substantially less support (P(M|data) = .134; BF_10_ = 0.347), providing moderate evidence against AQ as a predictor of d’. Adding AQ to the -model did not improve its predictive power compared to the model without AQ. An identical model comparison substituting AQ with the PCA-derived component yielded the same conclusion: the model including PCA (P(M|data) = .241; BF_10_ = 0.800) did not outperform the AQ-free baseline, providing converging evidence against the PCA component as a predictor of *d*’.

#### Autistic Traits Predict Decision Bias: PCA-modulated Anchoring

Having established that autistic traits do not predict perceptual sensitivity, we examined whether they instead modulate decision strategy or the relative weighting of standard-based versus positional anchoring. Because the standard-bias criterion reverses sign across standard positions by construction, collapsing across conditions would produce algebraic cancellation. We therefore modelled choice behavior at the trial level via GLMM, followed by condition-specific SDT criterion analyses.

#### Trial-Level Evidence: Generalized Linear Mixed Model

To test whether the PCA-derived component predicted first-stimulus anchoring, we fitted a trial-level GLMM with binomial family and logit link function. This approach leverages all ∼700 observations per participant and avoids the algebraic cancellation that can arise when collapsing across standard positions. Beyond the expected dominant main effect of STANDARD POSITION (χ^2^(1) = 97.53, *p* < .001), the model revealed a significant main effect of PCA (χ^2^(1) = 5.74, *p* = .017), indicating that higher PCA scores predict an increased probability of selecting the first-presented stimulus regardless of task duration or standard position (Figure 4). Crucially, the Task × PCA interaction did not reach significance (χ^2^(1) = 3.29, *p* = .070), providing no evidence that the anchoring tendency differed systematically between the Short and Long conditions. Similarly, the STANDARD POSITION × PCA interaction was non-significant (χ^2^(1) = 0.018, *p* = .893), confirming that first-stimulus anchoring in high-PCA individuals operated independently of whether the standard or comparison appeared first.

**Figure 4.**
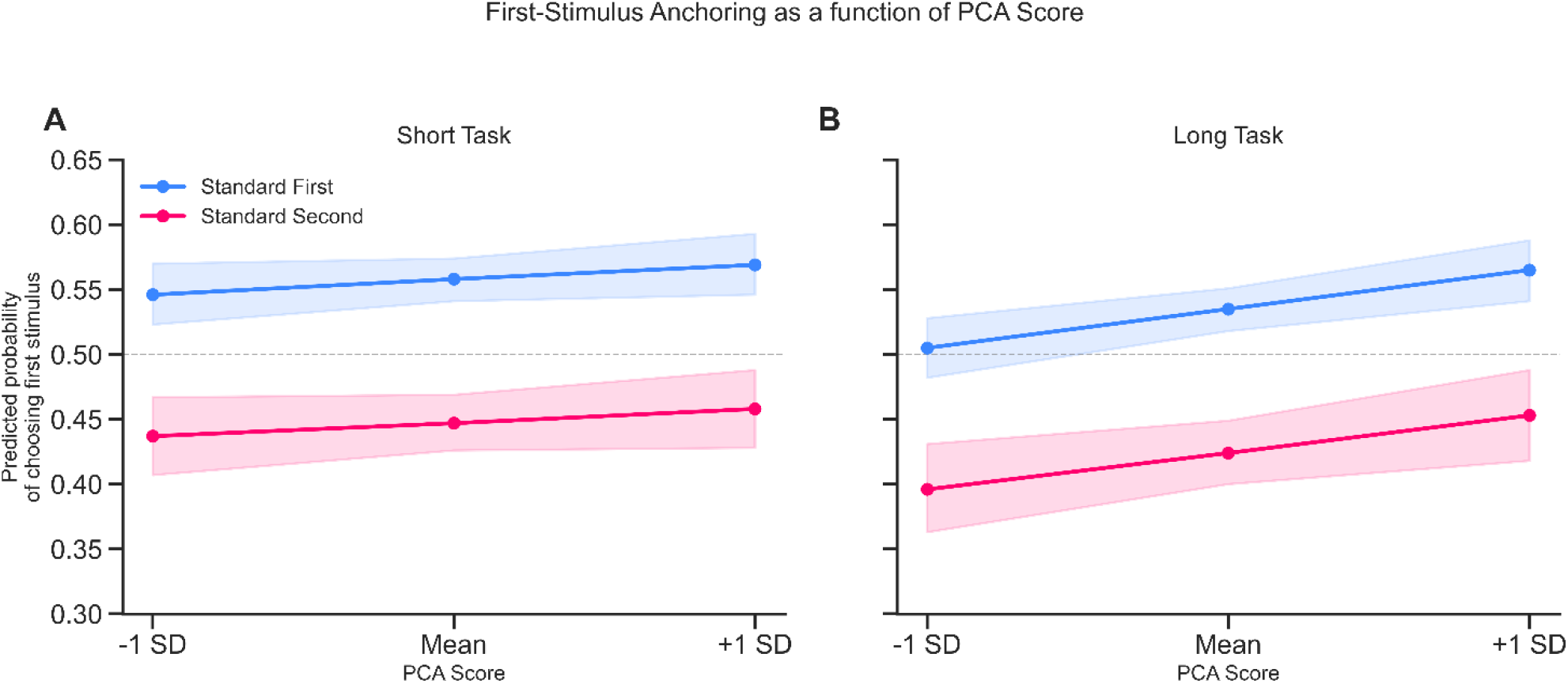
Predicted probability of first-stimulus selection as a function of PCA score, derived from the Generalized Linear Mixed Model. Panels A and B show the Short and Long tasks, respectively. The y-axis represents the predicted probability of selecting the first-presented stimulus; the x-axis represents the PCA component at −1 SD, Mean, and +1 SD. Shaded areas indicate 95% confidence intervals. The vertical offset between lines reflects the sample-level bias toward the standard:. when the standard appeared first (blue), participants selected the first stimulus more often than chance, as standard identity and ordinal position were aligned. When the standard appeared second (red), participants continued to favor it. Hence first-stimulus selection fell below chance, confirming that the sample as a whole anchored to the standard rather than to ordinal position. Critically, both lines show a positive slope, indicating that higher PCA scores increase first-stimulus selection regardless of standard-position, confirmed by the non-significant PCA × STANDARD POSITION interaction (χ^2^(1)=0.018, *p*=.893). The effect is graded: even at +1 SD, participants in the standard-second condition still favored the standard, but less strongly than lower-PCA individuals, reflecting a quantitative reduction in standard-based anchoring rather than a switch to positional heuristics.

This pattern indicates that higher PCA scores are associated with a generalized shift toward positional anchoring. Importantly, the absence of interaction effects demonstrates that this shift does not depend on whether the first stimulus is the standard or the comparison. Rather than selectively weakening standard-based anchoring in specific configurations, higher-PCA individuals show a stable increase in first-stimulus weighting across the entire task. The following SDT criterion analyses decompose the condition-specific structure of this effect.

### Condition-Specific Structure: SDT criterion analyses

#### Standard-bias criterion

Because its expected direction reverses across standard positions, the sign reversal in correlations is not a condition-specific effect but the algebraic expression of a single underlying tendency, consistent with the GLMM main effect. Spearman correlations between the PCA component and criterion values revealed significant associations in standard-first conditions across both tasks. In the Short Task, a negative correlation emerged in the standard-first condition (r=−0.33, *p*<.05), indicating that higher PCA scores were associated with stronger bias toward the standard when it appeared first; no significant association was found in the standard-second condition. In the Long Task, a comparable negative correlation was observed in the standard-first condition (r = −0.33, *p*<.05), alongside a significant positive correlation in the standard-second condition (r =0.29, *p*<.05). This sign reversal confirms that high-PCA individuals anchored to the first-presented stimulus rather than to the standard per se. AQ total scores showed a weaker pattern: only the Long Task standard-first condition reached significance (r = −0.25, *p* = .042), with no significant associations in the remaining conditions (Short Task: standard-first r= −.18, *p*=.146; standard-second r =.07, *p*=.556; Long Task: standard-second r=.14, *p*=.285), suggesting that the PCA component captures decision-relevant variance more selectively than the composite AQ score.

Linear regressions confirmed that PCA predicted criterion. In the short task, the effect was significant only in the standard-first condition (R^2^=0.096, F(1,63)=6.72, p=.012; β=−0.078), indicating stronger bias toward the standard at higher PCA scores; the standard-second condition was non-significant (R^2^=0.006, F(1,63)=0.38, p=.542; β=0.022). In the long task, both conditions were significant: standard-first showed a negative association (R^2^=0.111, F(1,63)=7.91, p=.007; β=−0.107), again reflecting standard-based anchoring; standard-second showed a positive association (R^2^=0.103, F(1,63)=7.20, p=.009; β=+0.114), consistent with positional anchoring toward the first-presented stimulus (the comparison).

Parallel regressions using AQ total score as predictor yielded no significant effects across any condition (Short Task standard-first: R^2^ =.016, F(1,63) = 0.994, *p*= .323; standard-second: R^2^ = .006, F(1,63) = 0.365, *p* = .548; Long Task standard-first: R^2^=.033, F(1,63) = 2.149, *p* =.148; standard-second: R^2^ = .016, F(1,63) = 1.027, *p*=.315), confirming that the PCA-derived component captures decision-relevant variance more selectively than the composite AQ score.

#### Order-bias criterion and positional anchoring across subsecond scales

To investigate whether PCA-related anchoring reflects a stable trait or a stimulus-specific phenomenon, we analyzed response criterion (c) as an index of first-versus second-stimulus preference (order bias), regardless of the standard’s identity.

In the Short Task, no significant association emerged when collapsing across positions (r =−0.17, *p* =.16). A significant correlation appeared in standard-first trials (r =−0.33, p=.008; Fig. 5A), indicating stronger first-stimulus bias at higher PCA scores, but not in standard-second trials (r = −0.09, *p*=.46; Fig. 5B). Rather than a genuine task-by-position dissociation, this pattern likely reflects a synergy of effects: in standard-first trials, the general tendency to anchor to the standard and the PCA-related bias toward the first interval operate in the same direction, whereas in standard-second trials, these tendencies compete, increasing noise at the aggregate level. Crucially, the GLMM, which leverages trial-by-trial power, confirmed a stable effect of PCA on response bias (*p* = .017) with no significant Task × PCA interaction (*p* = .070), suggesting that the underlying strategic shift is robust even when its expression is modulated by stimulus-specific constraints (see *Trial-Level Evidence: Generalized Linear Mixed Model* above).

**Figure 5.**
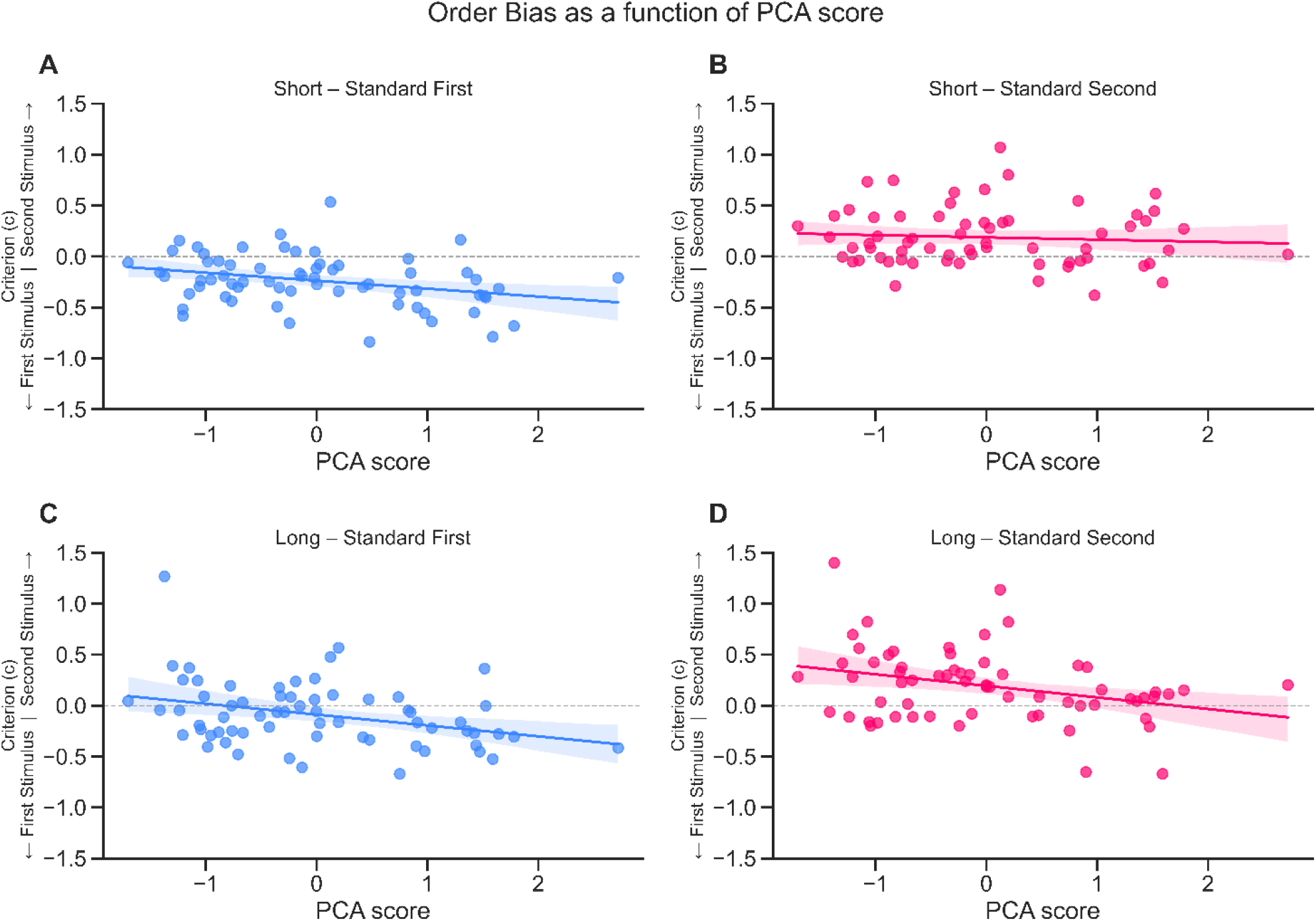
Order Bias as a function of PCA score across Task and Standard Position. Each panel shows the relationship between PCA score and the order-bias criterion (c). Negative values indicate a bias toward the first-presented stimulus; positive values indicate a bias toward the second. Regression lines with 95% confidence intervals are displayed. In the Short Task, a negative association with PCA scores was observed only when the standard appeared first (A), whereas no clear relationship emerged when it appeared second (B). In the Long Task, the negative association was present in both standard positions (C, D), suggesting a more consistent tendency to favor the first-presented stimulus.

In the Long Task, positional anchoring was consistent across all conditions: collapsed (r = −0.29, p<.05), standard-first (r = −0.33, p<.01), and standard-second (r =−0.29, p<.05). Skipped Spearman correlations, which account for bivariate outliers (Pernet, Wilcox, & Rousselet, 2012), confirmed robustness: collapsed (r =−0.26, 95% CI [−0.469, −0.036]), standard-first (r =−0.27, 95% CI [−0.50, −0.01]), standard-second (r =−0.27, 95% CI [−0.51,−0.02]). Collectively, these findings indicate that higher autistic traits are associated with a systematic response bias toward the first-presented stimulus, suggesting a decision strategy that prioritizes immediate temporal input over a stabilized internal reference built across trials.

## Discussion

Do individuals with higher autistic traits have an impaired perception of time? Using a Signal Detection Theory (SDT) approach, we distinguished perceptual sensitivity from decision bias in a sub-second temporal discrimination task. Results indicate that total autistic traits (AQ) are not associated with reduced sensitivity (*d*’), and Bayesian analyses provided converging moderate evidence against a perceptual deficit: neither AQ total scores nor the PCA-derived component improved model fit beyond task and position alone. These findings are consistent with prior studies reporting intact or superior subsecond discrimination in ASD or high-autistic-trait samples (Mostofsky et al., 2000; Jones et al., 2009; Falter et al., 2012, 2013), suggesting that perceptual encoding of duration magnitude is preserved along the autism-trait dimension.

More informative is what emerged at the level of decision strategies. At the sample level, the standard-position effect was large and robust, replicating a well-established finding consistent with the Internal Reference Model (IRM; Dyjas et al., 2012; Lapid et al., 2008): repeated exposure to the standard stabilizes an internal representational trace that anchors comparison judgments, facilitating discrimination when the standard appears first and degrading it when it appears second.

The theoretically decisive finding emerged when individual differences were examined. The PCA-component, combining heightened Attention to Detail and reduced Imagination, was systematically associated with a shift in anchoring strategy. In standard-first trials, higher PCA scores predicted stronger bias toward the first-presented stimulus; in standard-second trials, the direction reversed, predicting bias toward the first-presented comparison. This sign reversal indicates that individuals with higher autistic traits did not selectively anchor to the repeated standard but showed a generalized tendency to anchor to whichever stimulus appeared first. The GLMM confirmed this at the trial level: the main effect of PCA on first-stimulus selection was significant and stable across standard positions, with no reliable PCA × Standard Position interaction, indicating a generalized shift in cue weighting toward temporal order, independent of standard identity.

Typical observers exploit the repeated standard across trials, building a stabilized reference that overrides positional cues. Individuals higher on the PCA component appear to rely less on this standard-based anchor and more on within-trial ordinal position. Within the IRM framework, this is consistent with reduced effective weighting of accumulated cross-trial information during reference updating, resulting in a reference representation that is less stabilized by the statistical regularity of the repeated standard. The behavioral consequence is not impaired duration encoding but a redistribution of anchoring weight toward positional order.

The cognitive profile driving this dissociation is interpretively coherent. Reduced Imagination, reflecting diminished capacity to generate flexible internal representations and simulate non-present events, may limit the prior-building operation required to retain and generalize a temporal standard across trials. Heightened Attention to Detail indexes a processing style more responsive to immediate sensory input than to accumulated contextual structure. Together, this profile characterizes a system that privileges bottom-up, within-trial information over top-down, cross-trial representational knowledge.

This interpretation aligns with multiple theoretical accounts. Predictive coding frameworks (Pellicano & Burr, 2012; Van de Cruys et al., 2014) propose attenuated weighting of top-down priors in ASD (Tarasi et al., 2025); an internal reference model is precisely such a prior, and reduced exploitation of it manifests as diminished standard reliance and increased dependence on the first stimulus. Weak central coherence (Happè & Frith, 2006) predicts reduced integration of information across instances or the operation required to construct a stable cross-trial temporal average. Enhanced perceptual functioning accounts (Mottron et al., 2006) are also consistent: heightened sensitivity to immediate sensory detail coupled with reduced contextual integration produces the pattern observed here.

Autistic traits did not abolish standard-based anchoring; they attenuated it, shifting relative weight toward positional heuristics. This quantitative shift, rather than a categorical deficit, is consistent with a dimensional view of autistic traits and with the heterogeneity of temporal difficulties in ASD.

A unifying interpretation is that individuals higher in autistic traits show a reduced tendency to extract and exploit statistical regularities from temporal context. This context-dependent process appears less strongly engaged along the autism-trait continuum, not because perceptual encoding is impaired, but because reliance on cross-trial representational structure is reduced. The result is a shift toward a more rigid, context-independent strategy: anchoring to the first-presented stimulus, an order-based heuristic that requires no cross-trial updating. This aligns with Weak Central Coherence (WCC; Frith, 1989, 2003; Frith & Happé, 1994; Happé, 1999), which propose a reduced drive to integrate information into global representations.

These findings help reconcile the inconsistent literature. These findings offer a principled account of the inconsistent literature. The divergence between studies reporting intact or superior discrimination (Mostofsky et al., 2000; Jones et al., 2009; Falter et al., 2012, 2013) and those reporting elevated thresholds or greater variability (Bhatara et al., 2013; Kargas et al., 2015; Grondin, 2020) may reflect differences in task structure and anchoring demands rather than genuine perceptual inconsistency. Without separating d’ from criterion, what are in fact stable individual differences in representational strategy are easily misread as perceptual variation.

Clinically, temporal difficulties in ASD (e.g., anticipating transitions, adapting to temporal schedules) may reflect reduced spontaneous exploitation of statistical regularities rather than impaired duration perception, consistent with broader characterizations of autistic cognition including preference for predictable sequences and greater weighting of sensory-driven processing (Boucher et al., 2007; Tarasi et al., 2021; Pellicano & Burr, 2012; Van de Cruys et al., 2014).

## Limitations

Our sample reflects subclinical traits, consistent with a continuum view of autism. Findings are limited to sub-second visual intervals; future works should extend to longer durations and other modalities. The correlational design precludes causal inference, and future studies would benefit from additional behavioral or physiological indices alongside the AQ.

## Conclusion

Autistic traits are more strongly associated with shifts in decision strategy than with perceptual deficits in temporal discrimination. SDT revealed that first-stimulus anchoring is modulated by a profile of heightened Attention to Detail and reduced Imagination, suggesting interventions should target cognitive flexibility rather than perceptual remediation. Consistent with Gil et al. (2012), high-autistic-trait individuals possess the perceptual “raw material” for time estimation; differences in temporal behavior likely reflect higher-order processes rather than core perceptual deficits.

## Supporting information

Supplementary_PCA_PSE

## Acknowledgments

We thank Margherita Guandalini for help with data collection.

## Declaration of conflicting interest

None.

## Funding statement

VR is supported by Next Generation EU (NGEU) and funded by the Ministry of University and Research (MUR) through the National Recovery and Research Plan (NRRP), PRIN 2022 (grant no. 2022H4ZRSN — CUP J53D23008040006: Predictive waves in human perception and individual differences along the autism–schizophrenia continuum; grant no. P2022XAKXL — CUP J53D23017340001: Investigating the plasticity of human predictive coding through neuromodulation). Additional support was provided by the Ministerio de Ciencia, Innovación y Universidades, Spain (PID2019-111335 GA-100). SB and VR were also supported by the Bial Foundation (033/22).

## Ethical considerations

This study was approved by the Bioethics Committee of the University of Bologna (n. 0299334, 09/11/2022).

## Data availability statement

The data that support the findings of this study are available from the corresponding author upon reasonable request.

## Notes

### Competing Interest Statement

The authors have declared no competing interest.

