## Supplementary_PCA_PSE for "Autistic Traits are Associated with Suboptimal Decision Bias Strategies in Subsecond Timing"

**Supplementary Materials**

### S1. Principal Component Analysis of AQ Subscales

A Principal Component Analysis (PCA) was conducted on the five subscales of the Autism Spectrum Quotient (AQ): Social Skills, Communication, Attention Switching, Imagination, and Attention to Detail, using the correlation matrix as input.

Sampling adequacy was assessed using the Kaiser–Meyer–Olkin (KMO) measure and Bartlett's test of sphericity. The KMO index was acceptable (KMO = 0.650), and Bartlett's test of sphericity was significant ($\chi^2(10) = 42.84$, $p < .001$), confirming that the correlation matrix was suitable for component extraction.

Components were retained according to Kaiser's criterion (eigenvalues > 1) and inspection of the scree plot, which confirmed a two-component solution. The unrotated solution yielded two components with eigenvalues greater than 1: Component 1 (eigenvalue = 2.037, 40.7% of variance) and Component 2 (eigenvalue = 1.091, 21.8% of variance), together accounting for 62.6% of the total variance. To enhance interpretability, an orthogonal Varimax rotation with Kaiser normalization was applied (convergence in 3 iterations). After rotation, Component 1 explained 37.4% of the variance (rotated eigenvalue = 1.87) and Component 2 explained 25.1% (rotated eigenvalue = 1.26), with cumulative explained variance unchanged.

Component 1 (Social-Communication Dimension) showed high positive loadings for Attention Switching (.834), Social Skills (.767), and Communication (.691), reflecting social-interpersonal and cognitive flexibility difficulties characteristic of autistic traits. Component 2 (Detail–Imagination Dimension) showed high positive loadings for Imagination (.866) and Attention to Detail (.634), capturing a cognitive style characterized by heightened attention to local features and reduced imaginative flexibility. Higher scores on the Imagination subscale reflect reduced imaginative ability, while higher scores on Attention to Detail reflect an enhanced focus on local features. Individual component scores were extracted using the regression method and used as continuous predictors in subsequent analyses.

**Supplementary Table S1. Rotated Component Matrix (Varimax)**

| AQ Subscale | Component 1 | Component 2 | Uniqueness |
|---|---|---|---|
| Attention Switching | 0.834 | — | 0.291 |
| Social Skills | 0.767 | — | 0.347 |
| Communication | 0.691 | — | 0.494 |
| Imagination | — | 0.866 | 0.246 |

| AQ Subscale | Component 1 | Component 2 | Uniqueness |
|---|---|---|---|
| Attention to Detail | — | 0.634 | 0.494 |

***Note.*** *Extraction Method: Principal Component Analysis. Rotation Method: Varimax with Kaiser Normalization. Loadings < 0.30 suppressed. Uniqueness = 1 – Communality.*

**Supplementary Table S2. Variance Explained**

| Component | Eigenvalue (Unrotated) | % Variance | Cumulative % | Eigenvalue (Rotated) | % Variance (Rotated) |
|---|---|---|---|---|---|
| 1 | 2.037 | 40.7% | 40.7% | 1.87 | 37.4% |
| 2 | 1.091 | 21.8% | 62.6% | 1.26 | 25.1% |

***Note.*** *Component scores were extracted using the regression method.*

### S2. Point-Of-Subjective Equality Analyses

As an additional index of perceived duration, we estimated individual Points-of-Subjective-Equality (PSE) by fitting psychometric sigmoid functions to each participant's probability of judging the comparison interval as longer than the standard [$y = a + b/(1 + \exp(-(x - c)/d))$; Cecere, 2015]. The PSE was defined as the comparison duration corresponding to a response probability of 0.5, reflecting subjective equivalence between the two intervals (Block, Grondin, & Zakay, 2018; Grondin, 2024). Unlike d′ and criterion, the PSE does not dissociate perceptual sensitivity from decision bias and is reported here as a supplementary measure. Spearman correlations were computed between PSE values and both AQ total scores and the PCA-derived component for each task and standard position separately.

Collapsing across standard positions, the mean PSE was 106.4 ± 8.5 ms, with no significant correlations with the PCA component ($r = 0.09$, $p = 0.47$) or AQ total score ($r = -0.07$, $p = 0.58$). In the standard-first condition, five participants were excluded due to poor psychometric fits (adjusted $R^2 < 0.5$), yielding a mean PSE of 110.2 ± 19.2 ms. The correlation with the PCA component approached but did not reach significance ($r = 0.22$, $p = 0.08$), and no significant association was observed with AQ ($r = 0.17$, $p = 0.18$). No significant effects were observed in the standard-second condition.

In the standard-first condition, the mean PSE was 500.3 ± 86.9 ms, indicating a slight but significant underestimation of the comparison interval relative to the standard ($t(64) = 46.43$, $p < .001$, Cohen's $d = 5.76$). In this condition, PSE correlated positively with the PCA component ($r = 0.38$, $p < .005$) and AQ total score ($r = 0.27$, $p < .05$), indicating that higher scores on both the PCA component and AQ were associated with larger PSE values when the standard appeared first — a pattern consistent with an anchoring-related decision bias rather than altered duration encoding (N = 59, after excluding six participants with poor fits), the mean PSE was

536.2 ± 155.6 ms, with no significant correlations with the PCA component ($r = -0.19$, $p = 0.15$) or AQ ($r = -0.07$, $p = 0.58$). Collapsing across standard positions, the mean PSE was 515.1 ± 48 ms, again with no significant associations (PCA: $r = 0.08$, $p = 0.51$; AQ: $r = 0.13$, $p = 0.29$).

PSE effects were limited and condition-specific. The significant association observed in the Long Task standard-first condition is consistent with the anchoring account described in the main text. However, the absence of consistent effects across tasks and positions suggests that autistic traits do not produce a generalized alteration in perceived duration. In line with the SDT analyses, the overall pattern suggests that autistic traits modulate decision strategy rather than perceptual sensitivity.
